# Root Walker: an automated pipeline for large scale quantification of early root growth responses at high spatial and temporal resolution

**DOI:** 10.1101/2022.11.16.516796

**Authors:** Platre Matthieu Pierre, Mehta Preyanka, Halvorson Zachary, Ling Zhang, Brent Lukas, Gleason F. Matias, Faizi Kian, Goulding Callum, Busch Wolfgang

## Abstract

Plants are sessile organisms that constantly adapt to their changing environment. The root is exposed to numerous environmental signals ranging from nutrients and water to microbial molecular patterns. These signals can trigger distinct responses including the rapid increase or decrease of root growth. Consequently, using root growth as a readout for signal perception can help decipher which external cues are perceived by roots, and how these signals are integrated. To date, studies measuring root growth responses using large numbers of roots have been limited by a lack of high-throughput image acquisition, poor scalability of analytical methods, or low spatiotemporal resolution. Here, we developed the Root Walker pipeline, which uses automated microscopes to acquire time-series images of many roots exposed to controlled treatments with high-spatiotemporal-resolution, in conjunction with fast and automated image analysis software. We demonstrate the power of Root Walker by quantifying root growth rate responses at different time and throughput scales upon treatments with natural auxin, and upon treatment with two mitogen-associated protein kinase cascade inhibitors. We find a concentration-dependent root growth response to auxin and reveal the specificity of one MAPK inhibitor. We further demonstrate the ability of Root Walker for conducting genetic screens by performing a genome wide association study on 260 accessions under 2 weeks. This revealed known and unknown root growth regulators. Root Walker promises to be a useful toolkit for the plant science community, allowing large-scale screening of root growth dynamics for a variety of purposes, including genetic screens for root sensing and root growth response mechanisms.

## Introduction

The root is an organ that is involved in a plethora of functions, from anchoring the plant in the soil to facilitating nutrient and water uptake. However, resources are not evenly distributed in the soil. Moreover, plants interact with beneficial and pathogenic microorganisms which can promote growth, limit resources, or pose an immune threat. It is therefore not surprising that roots can sense and respond to many different stimuli, ranging from physical cues such as gravity and touch to chemical gradients such as water, nutrients or microbially-associated molecules (Morris *et al*., 2017). Root responses to the environment frequently result in a change in root growth dynamics. These responses are key for plant adaptation, and over time they shape the root system architecture (RSA). During the last decade, root growth phenotypes have been successfully used to identify novel genes and molecular mechanisms that underlie root responses to the environment (Slovak *et al*., 2014, 2020; Satbhai *et al*., 2017; Li, Sun, Huang, Goschl, *et al*., 2019; Platre *et al*., 2022). However, these studies screened relatively late readouts of root growth change, on the order of days after exposure to distinct growth environments. Such approaches might not be powerful enough to identify sensory mechanisms that are still unknown for many environmental cues such as nutrients (Jia, Giehl and von Wirén, 2022). We hypothesized that to capture these dynamics, it would be necessary to track root growth during the first hours after transfer to a treatment condition. However, existing tools to precisely measure such responses for hundreds or thousands of roots with high spatiotemporal resolution are sparse.

To accurately measure even small changes in root growth, spatial resolution is of high importance. While CCD scanners are otherwise suited for high-throughput studies (Slovak *et al*., 2014), they won’t yield the required resolution, as even high quality scans can only achieve ∼21 μm/pixel. Camera systems are frequently used to achieve medium resolution for root imaging and can detect subtle root growth changes (Durham Brooks, Miller and Spalding, 2010). Microscopes have been used in conjunction with growth chambers or microfluidic chambers to study root growth changes with high time resolution while inducing stress (Grossmann *et al*., 2011; Busch *et al*., 2012; de Luis Balaguer *et al*., 2016; von Wangenheim *et al*., 2017; Stanley *et al*., 2018). This approach is particularly effective at capturing rapid changes in root growth; however, the required hardware is expensive and can only track a small number of roots at a time, preventing high-throughput image acquisition. Moreover, the production of growth chambers, the use of confocal microscopy, and the lack of dedicated automation software suited for root biology frequently pose hurdles for their adoptability (Grossmann *et al*., 2011; Busch *et al*., 2012; de Luis Balaguer *et al*., 2016; von Wangenheim *et al*., 2017; Stanley *et al*., 2018).

Another important element required to study root growth in a high-throughput manner is the ability to automatically collect measurements at different timepoints. Many efforts have been made to develop root image analysis tools to meet this need. For instance, the MyRoot software package can measure root lengths at a high-throughput from scanner images (Betegón-Putze *et al*., 2019), but requires human interaction and thus limits the processing of longer time series. Tools that allow the assessment of root growth variation with high temporal resolution, such as RootTrace and RootTip (French *et al*., 2009; Geng *et al*., 2013), require the use of a camera which provides sufficient resolution to detect root growth variation on timescales from 3 minutes to 15 minutes, respectively. However, the use of a camera limits phenotyping to just a single plate of seedlings at a time. Moreover, the downstream image analysis is only semi-automated, which becomes increasingly time consuming with larger sample sizes. In other words, reliable tools that can acquire and automatically analyze root growth in high-throughput and with high spatiotemporal resolution are not currently available.

Here, we present Root Walker: an analytic phenotyping platform that enables the high-throughput acquisition and quantification of early root growth responses at high temporal and spatial resolution. Root Walker permits the acquisition of bright-field root images using a microscope beginning minutes after the application of a treatment and continuing for up to 12 hours. The acquired images are then processed using a MATLAB^®^ script in which roots are segmented and their growth after transfer is automatically quantified. To illustrate the accuracy and power of this method, we used *Arabidopsis thaliana* roots treated with different concentrations of the auxin analog indole-3-acetic acid (IAA), as well as two mitogen-associated protein kinase (MAPK) inhibitors (U0126 and PD98059). Additionally, to demonstrate the benefits of this platform for facilitating genetic screens that lead to the identification of new molecular players involved in root growth regulation, we performed a genome wide association study (GWAS) using 260 accessions. This screen was completed in less than 10 days. Using these GWAS data, we identified already known and unknown root growth regulators such as the *SUPPRESSOR OF NPR1-1* (*SNI1*) and *ETHYLENE RESPONSIVE ELEMENT BINDING FACTOR 1* (*ERF-1*). Moreover, we discovered that the maintenance of the anionic lipids phosphatidylinositol-4-phosphate (PI4P) levels by PI4 Kinases participates in regulating root growth.

## Results

### A simple growth chamber in conjunction with automated microscopy allows for root imaging at high temporal and spatial resolution

We set out to develop an automated platform for capturing root growth responses at large scale. For this, we wanted to image at a resolution that could capture even small changes in root growth. We also needed an automated platform to conveniently capture multiple roots at high temporal resolution. We used an inverted BZ-X810 KEYENCE^®^ microscope, which can automatically capture images at a resolution of ∼3μm/pixel using a 2X objective (NA/0.1) or ∼1.5 μm/pixel using a 4X objective (NA/0.2). This level of resolution allows the detection of subtle root growth rate changes. For capturing many roots at a time, we used an agar-filled chamber in which we could place ∼20 seedlings (Figure S1a). To allow the accurate quantification of root growth responses, plants were first germinated under standard growth conditions on plates before being transferred to chambers containing the treatment medium prior to imaging (Figure S1). For each chamber, either 9 or 17 images were acquired depending on the objective used. Three of these chambers can be mounted at a time (Figure S1b-c), resulting in an imaging capacity of ∼60 seedlings per experimental round. As we used 4 microscopes total, this further scaled to ∼240 roots at a time. The described imaging setup allows the capture of images at a fast rate which varies according to the objective used and the number of imaging chambers loaded on the microscope stage. This ranges from 18 seconds (for one chamber using the 2X objective) to 59 seconds (for three chambers with the 4X objective). To contend with these differences, we used two different setups, acquiring images once per minute for 60 minutes on the 4X objective, and once every 5 minutes for 12 hours with the 2X objective. Taken together, this hardware setup provides a platform to acquire images of roots at high temporal and spatial resolution in a high-throughput manner.

### A user-friendly software-tool extracts root lengths in an automated and high-throughput manner

After successful image acquisition, each image is processed by a user-friendly MATLAB^®^ script that only requires three user inputs: i) selecting a parent folder containing the images; ii) selecting a target folder to store the output files; and iii) supplying a descriptive prefix to be used for the output files. This script automatically quantifies *Arabidopsis thaliana* root lengths in the time series images acquired with the Keyence^®^ BZ-X810 microscope by performing several steps: (1) converting raw images to a large stitched image, (2) binarizing the stitched image, (3) converting the result into arrays specifying the midline of each root center, (4) “walking” along the midlines to virtually reconstruct each root, (5) calculating root lengths and removing outliers, (6) allowing quality control by generating videos and graphical plot output files, and(7) exporting the raw data in .csv files for (8) downstream statistical analysis (Figure 1a). The script is extremely fast, requiring only ∼190 ms to process a single image (or ∼1.710 seconds for a typical 9-image stitch) using a desktop computer with an Intel^®^ Xeon^®^ W-2135 CPU @3.70 GHz and 64GB of RAM. Note that we developed two different scripts adjusted to quantify root growth when acquiring images every minute for 60 minutes using the 4X objective, or every 5 minutes for 12 hours using the 2X objective, as mentioned in the previous paragraph. In the subsequent sections, we describe the software processing steps in detail.

**Figure 1.**
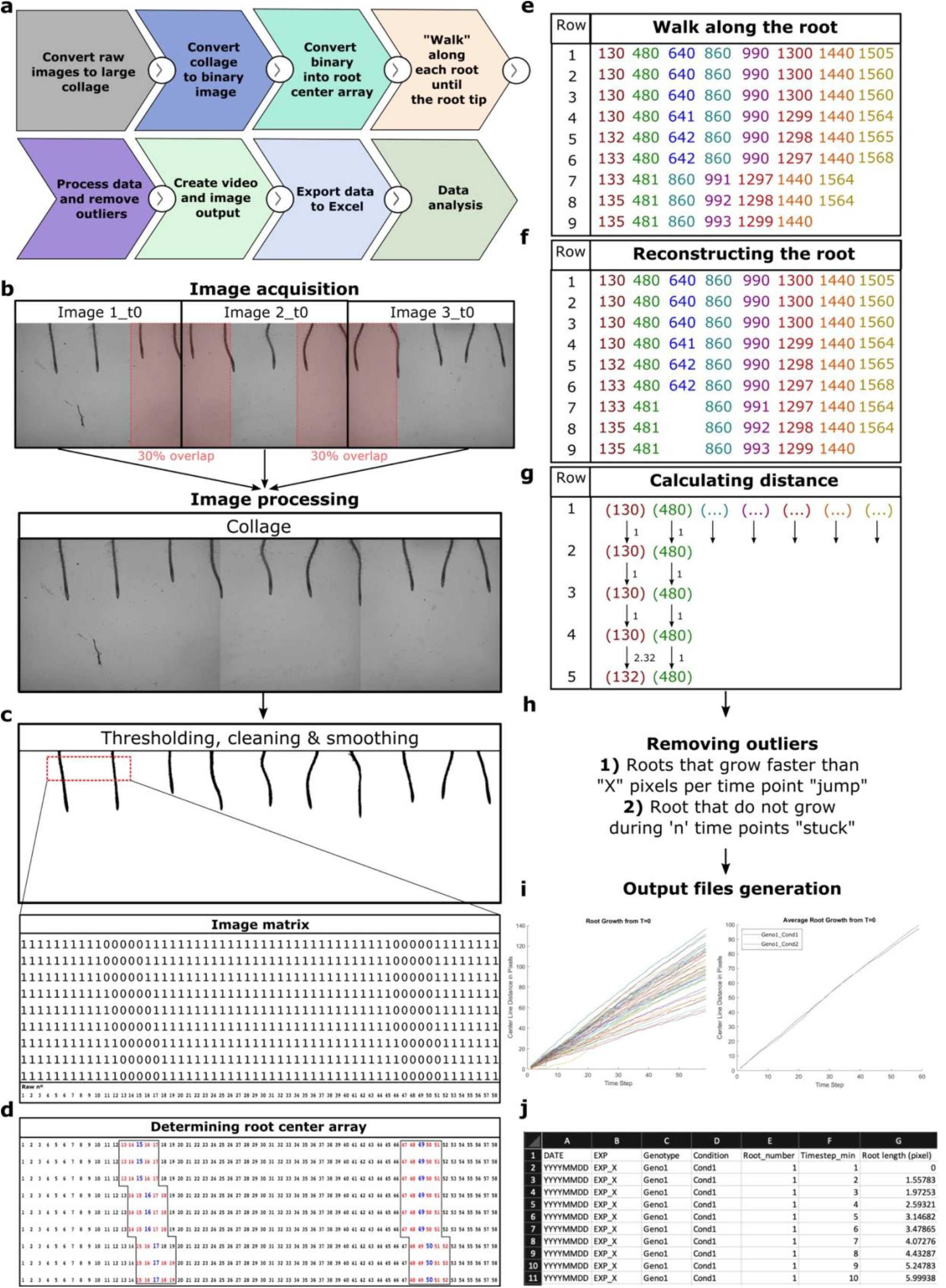
Overview of the Root Walker analysis platform. **(a**) Step diagram of the root Walker analysis platform. b) Pictures that are acquired overlap by 30%. (**b**) Stitched picture (collage). (**c**) Upper panel, binary image obtained by using three steps: thresholding, removing background and smoothing. Lower panel, binary matrix (subset from red box) obtained from binary image, 0 represents a black pixel and 1 a white pixel. (**d**) Determination of the Root Center Array from the binary image using connected sets of black pixels and extraction of the root center values (blue text). (**e**) Root center value array stored as a data sheet. The algorithm walks down each row in the matrix to determine if there is a connected object in the next row progressing vertically down the image. The connected objects are represented by the same color and the number represent the root center array value. (**f**) The connected objects are stored separately as arrays in a data sheet. (**g**) Distances are then calculated for each root array in the matrix to calculate the root length. (**h**) removal of improbable root growth values. (**i**) Graphical output file representing the root length (in pixels) of each root, left, or in for the same genotype in one condition according to time. (**j**) Example of the data stored in the .csv file when opened in Excel^®^.

### Root segmentation

The first step (1) of the algorithm consists of stitching all images that were acquired during a single time point together (Figure 1b). The next step (2) is to accurately segment all roots that are contained in the stitched image, which will enable the precise measurement of root lengths (Figure 1c). For this, a binarized image is generated using Otsu’s thresholding method, which minimizes the intraclass variance of the binary pixels (Figure 1c). Locally adaptative thresholding and darker foreground than background methods are applied to increase thresholding accuracy (Figure 1c). Finally, background noise and air bubbles coming from the camera sensor itself or from the plant transfer procedure to the imaging chamber are eliminated, respectively. This step removes small objects (< 8000 pixels) and very large objects (> 150 000 pixels) to obtain images with only roots as the remaining objects (Figure 1c). The resulting image contains only segmented roots. However, these segmented roots have jagged, fractal-like edges. If used to track the center line of each root directly, these lead to an overestimation of root length (analogous to the coastline paradox). To reduce this error, the image is convolved with a smoothing kernel to remove the jagged root edges. The smoothing kernel corresponds to a 3D average smoothing kernel with a window size of 21 that is set to return the output as the same size as the input image. The borders are then cleared of any black pixels created during the smoothing process (Figure 1c). The binarized stitched images are now ready for the subsequent generation of diagnostic videos and data analysis.

### Determination of root length using Root Walker

Once the images are segmented, (3) the root midline is determined for each root object across all time points (Figure 1d). This process consists of several steps: (1) for each row of each stitched image, the indices of the black pixels (Figure 1d) corresponding to an object are recorded. This set of indices is saved as an *n*-dimensional vector for each row of each stitched image, where *n* is the height of the image in pixels (Figure 1c, e.g. 1920 pixels). (2) The set of black pixel indices is separated into vectors of connected sets of black pixel indices, separating each object that is not touching or connected in the image. The objects are now considered distinct, which allows them to be processed individually. (3) The mean value of each vector of the black pixel indices is chosen to correspond to the center point index for each object. This greatly reduces the width of the dataset, reducing the computational cost of subsequent analysis (Figure 1e). This process is performed for each row of each stitched image.

The center point index array does not reflect the root length after transfer yet as each root object is stored in a data set which is the same height but for which the width is variable as indicated in Figure 1e. Therefore, further processing needs to be conducted by “walking” along each root pixel array to artificially reconstruct the root (step 4, Figure 1f). The value of the center point index array is used to determine which objects correspond to roots based on its horizontal distance in the next row. We step down each row from top to bottom and search for an index in the next row that lies within a maximum horizontal distance of the previous row’s center point. This parameter can be easily tuned by the user to obtain the best results on their dataset. By performing this process iteratively (Figure 1f) for each timestep, stitched image, starting point in the center point index array, and row of the stitched image, we can reconstruct every root. Our results now include position vectors for the midlines of every root in the image.

To increase the robustness of our root growth rate measurements, the software next integrates different timesteps and conducts further processing. We aimed to address curvature due to circumnutating (Taylor *et al*., 2021) that may introduce errors into the growth measurements, as well as potential distortions due to our focus on pixel-based midlines caused by data sparsity. To overcome these issues, only new growth is measured at each timepoint. This is achieved by overwriting each timestep with the previous one, ensuring that only new growth relative to the previous root tip location is measured. Next, the position vectors for each root are used to perform a midline distance calculation to determine the length of each root at each timestep (Figure 1g and Figure S2).

### Outlier removal

We also automatically reject growth rates that likely correspond to image processing errors, such as those caused by non-root objects being incompletely removed earlier in the method (step 5, Figure 1h). We do this by excluding any elements that do not change in length for a chosen number of timesteps; again, this threshold can be easily tuned by the user depending on their experimental setup. These outliers are often roots that are not growing due to damage incurred while transferring them to the imaging chamber (which we call “stuck” roots) or a non-root object (such as an agar pit or a bubble) that was included in the binarized image. We also remove any roots that “jump” and exhibit an unusually large length increase over a single timestep. We set this threshold at approximately ∼28μm per minute, which is over 7 times faster than normal root growth rates observed in healthy roots (∼3.9μm/min, Figure 1h). However, this parameter is also user-accessible to permit maximum flexibility. Depending on our experimental setup, “stuck” and “jump” events led to a loss of 8% of roots in the 60-minute time course (114 of 124 roots detected over 7 independent experiments, with 10 excluded) and 25% for 12-hour time course (129 of 171 roots detected over 6 independent experiments, with 42 excluded). We did not encounter any false positives. After removing outliers and artifacts, the root length for each timestep is used to determine the amount of new root growth per timestep. This is performed iteratively at each timestep to extract the complete root growth dynamics.

### Generation of output files

To allow for a quality control step in the analysis, our script also generates a timelapse video with annotations suitable for manual inspection (step 6, Video S1). Raw and processed stitched images are displayed side-by-side as a video overlaid with the extracted root midlines (Video S1). “Jump” and “Stuck” events are also shown. This step takes approximately half of the full image analysis time (∼95 ms per single image, or 855ms for a typical 9-image stitch). Skipping the video production step therefore leads to a significant improvement in processing speed.

To provide a succinct overview of the results, two plots are automatically generated (Step 6, Figure 1i). The first contains root lengths versus time, showing each individual root throughout the entire experiment. The second groups the roots by genotype and condition and plots the average root lengths for each group versus time. Both plots are saved and exported as .jpeg images. Reviewing them can help rapidly identify potential outliers that were not detected by the automated analysis, or aid users in tuning the script’s parameters for optimal results.

The processed root datasets are exported as a tidy .csv file (using pixels as units) to allow for comprehensive data analysis (Step 7, Figure 1j). This includes the date, replicate, genotype, and growth condition metadata, which are extracted from the parent folder. Pixel units can be converted into physical measurements with a simple scaling factor, and any root length derived traits, such as root growth rate and other parameters can be calculated (Step 8).

### Validation of the platform by studying early root growth responses to IAA

To evaluate the accuracy of our method, we compared our script’s root length quantification results to the measurements of trained human observers. No significant differences were observed between the two methods, showing that our tool produces accurate results (Figure S3a-b). The observed discrepancies between the script and any of the three human observers were not more profound than the differences between the human observers (Figure S3c-e). Overall, this shows that our method is as accurate as trained human observers. Furthermore, root growth rates extracted by our system were ∼2.3 +/-0.52 μm/min, which is in line with previous reports (Figure S3d; Shih *et al*., 2015; Fendrych *et al*., 2018; Prigge *et al*., 2020; Li *et al*., 2021; Serre *et al*., 2021; Dubey *et al*., 2023).

We next tested the utility of the Root Walker platform by studying the response of *Arabidopsis* roots to treatment with indole-3-acetic acid (IAA). To our knowledge this is the only treatment which has been reported to affect root growth within minutes when applied at nanomolar concentrations (Shih *et al*., 2015; Fendrych *et al*., 2018; Prigge *et al*., 2020; Li *et al*., 2021; Serre *et al*., 2021; Dubey *et al*., 2023). We applied IAA at 1nM (IAA[1]), 10nM (IAA[10]) and 100nM (IAA[100]), then tracked root growth by capturing images every minute for one hour. A standard condition control displayed root growth rates in line with our previous results (∼3.9+/-1.37 μm/min) (Figure 2). In accordance with the literature, we observed a dose-dependent decrease in the root growth rates in the treatment conditions, on the order of μm per minute (Figure 2 and Video S2, Serre *et al*. 2021). Within just 1 minute of transfer, a significant decrease of the root was observed for treatment with 10nM IAA and 100nM IAA (Figure 2b). At the lower concentration of 1nM IAA, the first significant difference of root growth rate was observed after 11 minutes (Figure 2b). These results highlight the Root Walker platform’s capability for capturing extremely fast root growth changes with high throughput and spatiotemporal resolution, making it a powerful tool for studying root growth responses.

**Figure 2.**
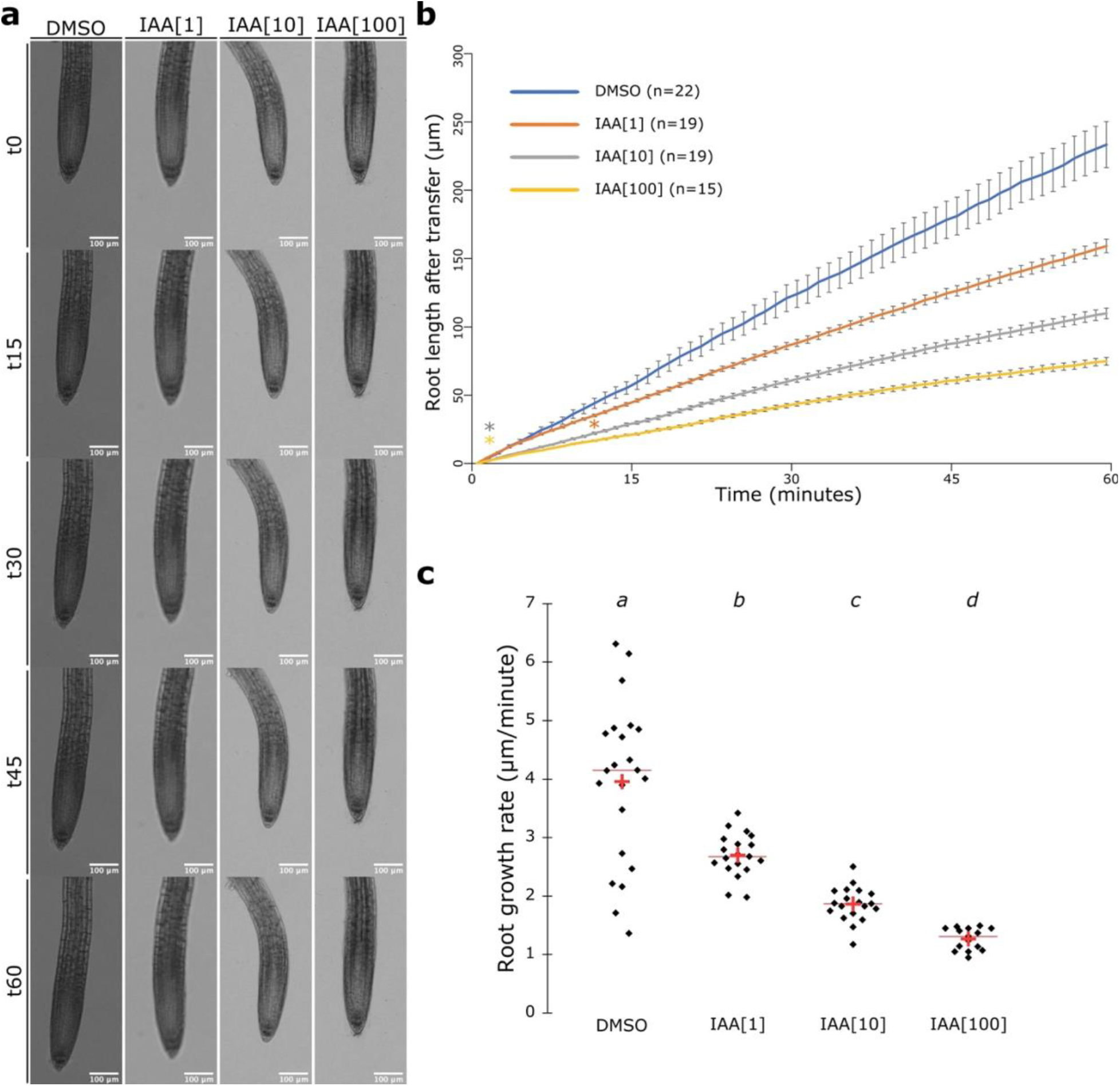
Exogenous application of indole-3-acetic acid for an hour restricts root growth in a concentration-dependent manner. (**a**) From left to right, representative images of roots of 5-day old seedlings treated with DMSO, IAA at 1nM, 10nM and 100nM for over the course of one hour. Scale bar, 100 μm. (**b**) Graphical representation of the root length increases over time after transfer to treatment medium. [two-way student t-test; (p<0.05)]. Asterisks indicate statistical differences compared to the DMSO and the color corresponds to the curve. Error bars represent the standard deviation. (**c**) Quantification of root growth rate expressed in μm per minute of seedlings treated with DMSO, IAA at 1nM, 10nM and 100nM for an hour. [one-way ANOVA followed by a Fisher LSD test; letters indicate statistical differences (p<0.05)]. Red crosses represent the average and the red bars the median. Black dots individual measurements

### Investigation of the effect of two MAPK inhibitors on root growth over 12 hours

After demonstrating that our platform is well-suited to studying rapid responses, such as that to IAA, we also wanted to test whether it was capable of accurately capturing growth responses that only occur after several hours. For this, we investigated the impact of two well-known mitogen associated protein kinase homolog 2 (MPK2) inhibitors on root growth. The ubiquitous MPK2 is part of the MAPK phosphorylation cascade, which is central in the regulation of eukaryotic developmental signaling events (Ge, Fu and Meadows, 2002; Hotokezaka *et al*., 2002; Bjornson *et al*., 2014). We used two inhibitors of MPK2 that have different modes of action: PD98059, which inhibits the MAPK cascade by binding the inactive form of MPK2, and U0126, which directly inhibits MPK2 kinase activity (Bjornson *et al*., 2014). To study the root growth response to these MPK2 inhibitors, we monitored root length for 12 hours, taking images every 5 minutes. We noticed that the root growth rate (mm/hour) was significantly lower when plants were exposed to U0126, but no differences were apparent for PD98059 compared to the mock treatment (Figure 3 and Video S3). Moreover, a statistically significant difference in root length between U0126 and mock conditions was already apparent after 2 hour and 25 minutes (Figure 3b). We concluded that the inhibition of MPK2 kinase activity, rather than the sequestration of the inactive form, is critical in regulating root growth. Our experiments demonstrate that the Root Walker platform can shed light on new regulatory mechanisms involved in early root growth.

**Figure 3.**
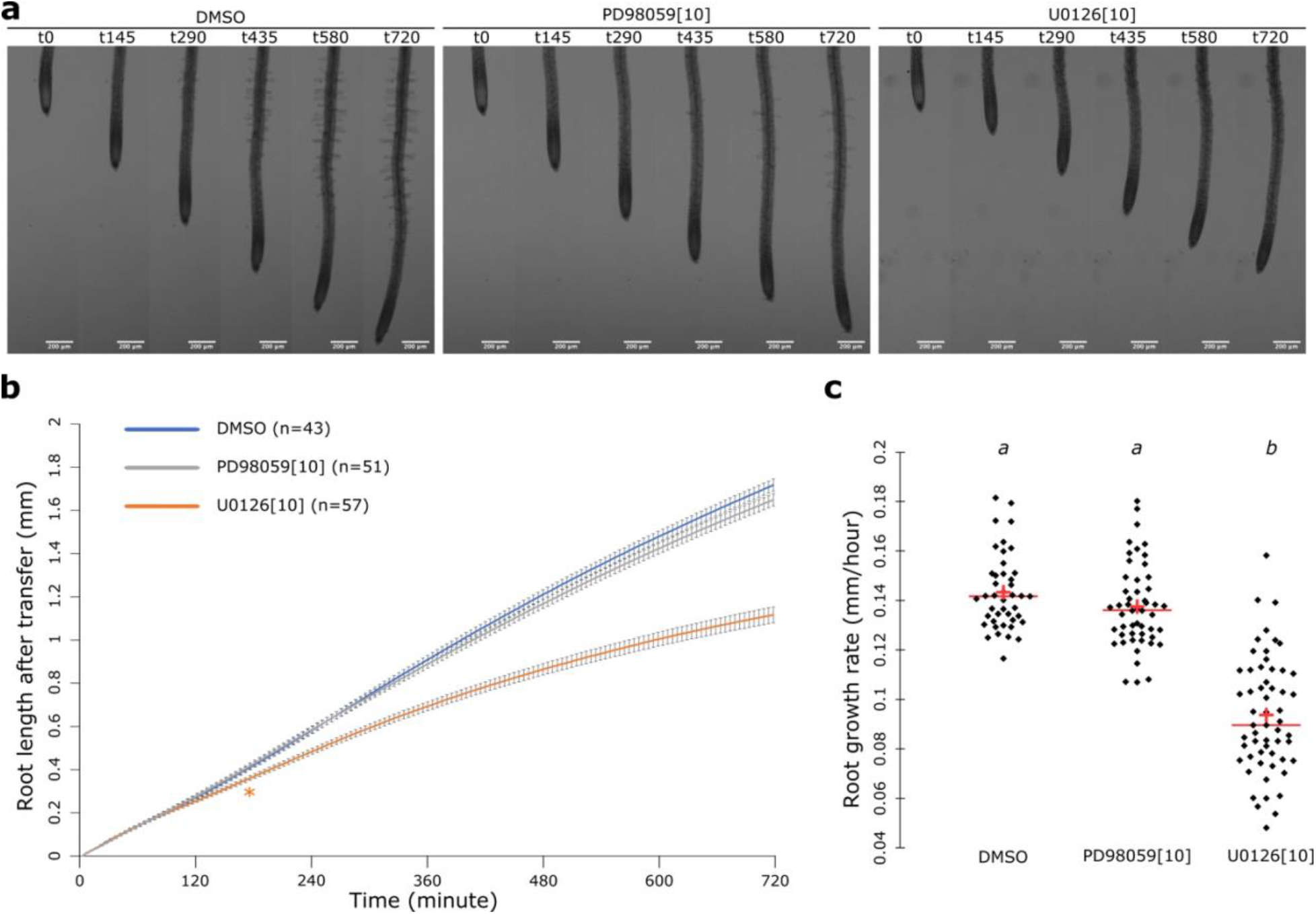
Exogenous application of the MPK cascade inhibitor U0126 but not PD98059 restricts root growth. (**a**) From left to right, representative images of roots of 5-day old seedlings treated with DMSO, PD98059 and U0126 at 10μM over the course of 12 hours. Scale bar, 200 μm. (**b**) Graphical representation of the root length increases after transfer to DMSO, PD98059 and U0126 at 10μM for 12 hours. [two-way student t-test; (p<0.05)]. Asterisks indicate statistical differences compared to DMSO and the color corresponds to the curve. Error bars represent the standard deviation. (**c**) Quantification of the root growth rate expressed in mm per hour of seedlings treated with DMSO, PD98059 and U0126 at 10μM for 12 hours. [one-way ANOVA followed by a post hoc Tukey HSD test; letters indicate statistical differences (p<0.05)]. Red crosses represent the average and the red bars the median. Black dots individual measurements.

### Assessing natural variation of early root growth responses for conducting genome wide association studies

Large-scale phenotyping enables quantitative genetic approaches such as genome wide association studies (GWAS) that can lead to the identification of new genes and molecular mechanisms for root growth (Satbhai *et al*., 2017; Li, Sun, Huang, Göschl, *et al*., 2019; Ogura *et al*., 2019; Slovak *et al*., 2020; Platre *et al*., 2022). While fast root growth responses had yet not been accessible for such approaches as they require the phenotyping of hundreds of lines, we were able to phenotype root growth of 260 accessions (Table S1) using our pipeline in less than two weeks. We used a condition (1/2MS, 0.5% sucrose and 0.8% agar with 100nM EtOH) that would be suitable as a control treatment for studying growth responses to many plants hormone, chemical, or small bioactive molecule. We then assessed multiple traits such as the mean, median and variance of the root growth rate in μm per minute along with the area under the curve per accession. We found notable variation of these root growth traits between the accessions in our phenotyping condition (Figure 4). This variation was heritable according to the broad sense heritability (BSH), which ranged from 37% to 61% (Figure 4). The observed BSH was in line with BSH observed for root traits from plants grown on agar plates (Satbhai *et al*., 2017; Li, Sun, Huang, Göschl, *et al*., 2019; Ogura *et al*., 2019; Slovak *et al*., 2020; Platre *et al*., 2022,). We then performed a mixed model based GWAS approach that corrects for population structure and detected a total of 389 SNP associations that passed the Benjamini Hochberg threshold of 5% (Figure 4, S4 and Table S2). The mean root growth rate led to the identification of 139 loci, the median root growth rate to 104 loci, the variance of root growth rate to 99 loci, and the area under the curve to 47 loci. Some loci were shared between different traits resulting in 345 unique loci. Considering a window of 20Kb around each identified loci, we identified 430 genes. We then selected the 64 genes that were associated with at least two different traits (Table S2) and determined in which specific root cell types these cells were expressed using COPILOT (Hsu *et al*., 2022). This revealed an enrichment of associated genes in the quiescent center and meta phloem & companion-related tissues (Figure 4e,).

**Figure 4:**
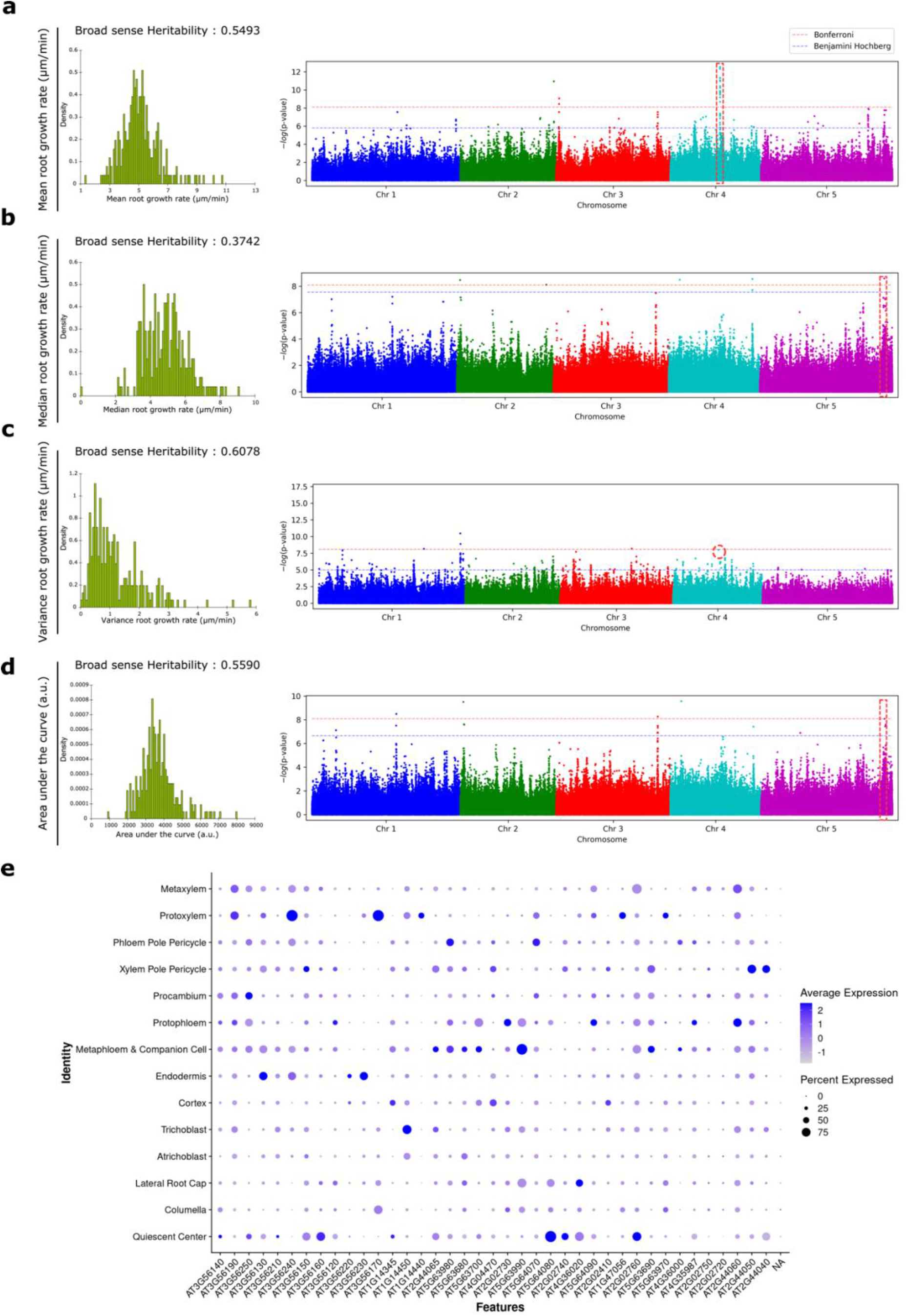
GWAS on root growth identifies multiple candidate genes. (**a-d**) From left to the right, broad sense heritability value (top), histogram (bottom) and Manhattan plot of mean root growth rate (**a**), median root growth rate (**b**), variance of root growth rate (**c**), area under the curve (**d**). (**e**) Dotplot of the genes associated with loci that were significantly associated with at least two of the measured root growth associated traits. NA stands for the genes from where the single cell data was not available. Dot size represents the percentage of cells in which each gene is expressed (% expressed). Dot colors indicate the average scaled expression of each gene in each cell-type group with warmer colors indicating higher expression levels.

To further validate our data set, we focused on a subset of associated loci and genes. The most significantly associated SNP for mean root growth rate (Chromosome 4, position: 10193934, p-value = 3.00E-13) was detected as being positioned in the coding region of AT4G18470 (Figure 4a). Moreover, this gene has the lowest relative distance from the related SNP (1025pb) compared to any other genes in the 20 kb window (1025bp, Table S2). It encodes for the *SUPPRESSOR OF NPR1-1* (SNI1,) for which the mutant has been described with significant shorter root (Wang *et al*., 2018). Moreover, we observed loci that were significantly associated with different traits. One SNP that was highly significantly associated with median root growth rate and area under the curve (Chromosome 5, position: 25645925, p-value= 2.51E-08) was 8949pb upstream from *PHOSPHATIDYLINOSITOL 4 PHOSPHATE BETA 1* (*PI4Kβ1*) and 2088 downstream from *HYCCIN2*. Notably, both genes had been shown to be involved in the biosynthesis of the PI4P (Okazaki, Miyagishima and Wada, 2015; Noack *et al*., 2021; Gomez *et al*., 2022). This prompted us to test the involvement of PI4K in root growth regulation by inhibiting its activity using the specific and well characterized small molecule, phenylarsine oxide (PAO, Simon *et al*., 2016; Platre *et al*., 2018). Consistent with our hypothesis that one or two of these genes playing a role in the determination of root growth rate, we observed that PAO caused a decrease of root growth in a dose dependent manner (Figure 5a-b and Video S4). Importantly PAO decreased the amount of cellular PI4P within this time frame showing that sustaining a constant PI4P pool at the plasma membrane through PI4Ks is partly involved in determining root growth (Figure 5c). Finally, we analyzed a significant associated locus emanating from the GWAS performed on the variance trait. The significantly associated SNP (position: 9761910, p-value= 6.73E-07) is in proximity of two genes AT4G17500 and ATG4G17505 corresponding to *ETHYLENE RESPONSIVE ELEMENT BINDING FACTOR 1* (*ERF-1*) *and PROTEIN OF UNKNOWN FUNCTION 239* (*DUF239*), respectively. To determine which of these genes might be involved in determining root growth, we took advantage of Transplanta lines, which contain inducible over-expresses specific transcription factors (Coego *et al*., 2014). We found that all three Transplanta lines showed a significant decrease in root growth after 3 days of induction that was similar to the positive control TOR-ES#1 and #2 which displayed a relatively small root upon induction due to defect in root meristem cell maintenance (Xiong and Sheen, 2012; Xiong *et al*., 2013; Figure 6 and Video S5). These results suggest that the transcription factor ERF-1 might be involved in root growth regulation.

**Figure 5.**
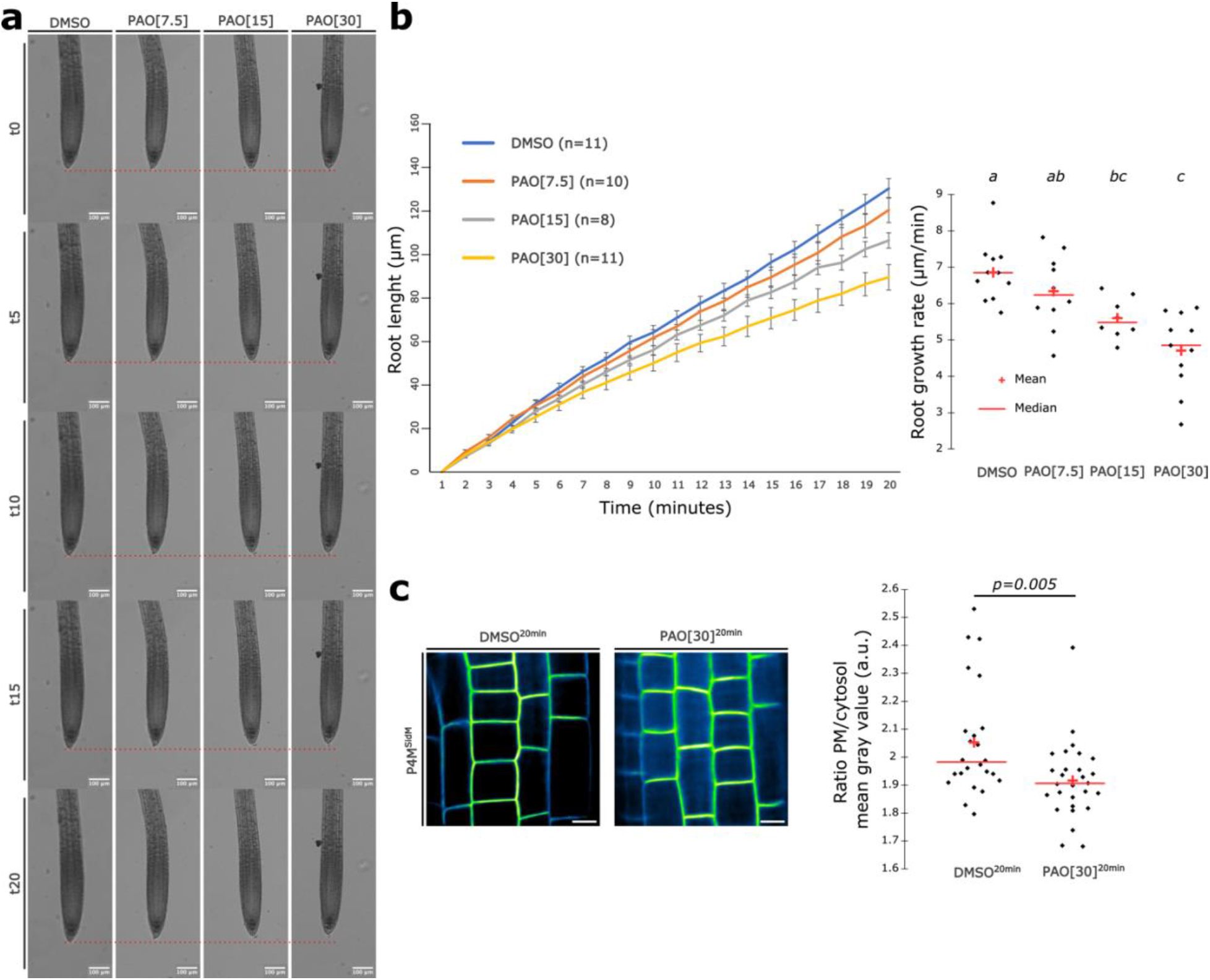
Exogenous application of the PI4 kinase inhibitor PAO restricts root growth in a concentration dependent manner. (**a**) From left to right, representative images of roots of 5-day old seedlings treated with DMSO, PAO 7.5μM, PAO 15μM and PAO 30μM over the course of 20 minutes. Scale bar,100 μm. (**b**) Graphical representation of the root length increases over time after transfer to treatment medium. Error bars represent the standard deviation and the related quantification of root growth rate expressed in μm per minute of seedlings treated with DMSO, PAO 7.5μM, PAO 15μM and PAO 30μM over the course of 20 minutes. [one-way ANOVA followed by a Fisher LSD test; letters indicate statistical differences (p<0.05)]. Red crosses represent the average and the red bars the median. Black dots individual measurements. (**c**) Confocal images of root epidermal cells of 5 day old seedlings expressing *pUBQ10::mCITRINE-P4M*^*SidM*^ with DMSO and PAO 30μM for 20 minutes and the related quantification. [two-way student t-test; (p<0.05)]. Red crosses represent the average and the red bars the median. Black dots individual measurements.

**Figure 6.**
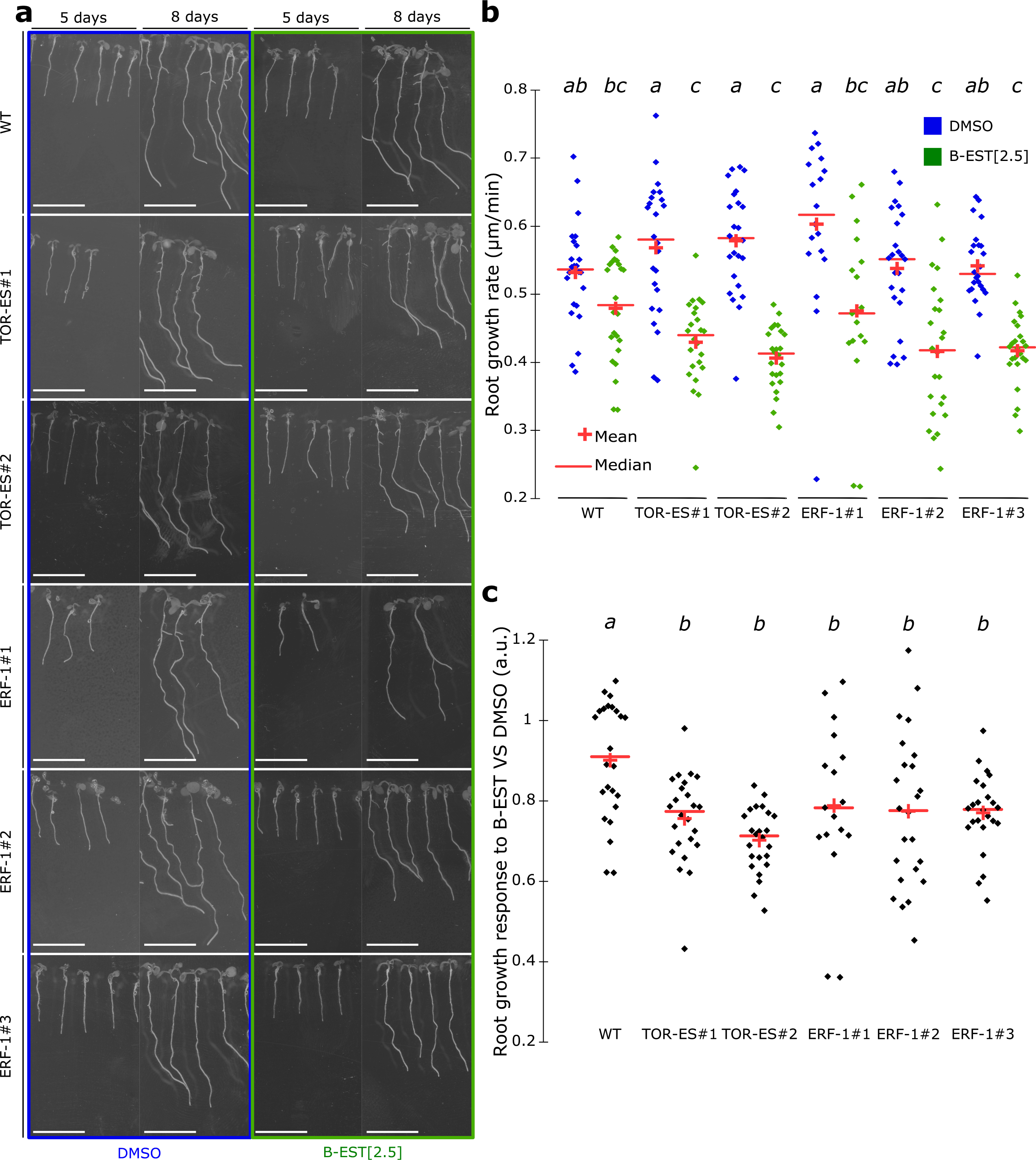
Inducible over expression of the transcription factor ERF-1 impairs root growth. (**a**) Left in blue rectangle, images of 5- and 8-days old Arabidopsis seedlings in DMSO, right in green rectangle, images of 5- and 8-days old Arabidopsis seedlings in β-estradiol (β-EST) at 2.5μM of WT, TOR-ES#1, TOR-ES#2, ERF-1#1, ERF-1#2, ERF-1#3. Scale bar, 1 cm. Note that TOR-ES lines is a conditional TOR mutant that has been shown to present significant root growth reduction upon induction. (**b-c**) Related quantification of the root growth rate in mm per day, (**b**) and the root growth response to in β-estradiol (**c**) [one-way ANOVA followed by a Fisher LSD test; letters indicate statistical differences (p<0.05)]. Red crosses represent the average and the red bars the median. Black dots individual measurements.

Taken together, these results clearly show the power of the Root Walker pipeline to expose new root growth regulatory mechanisms using genetic screens that can be conducted faster than most other available methods.

## DISCUSSION

Here, we have described a new high-throughput platform that can accurately quantify early root growth responses to stimuli with high spatiotemporal resolution. Roots can be imaged just minutes after transferring them to media containing a treatment condition, thanks to the use of agar-based imaging chambers. Each chamber can accommodate around 20 roots. By using 4 automated digital microscopes, each of which holds up to 3 chambers, we were able to image a total of 240 roots in parallel. Using such a system, a time resolution of more than 30 seconds after the transfer could be achieved at a spatial resolution of ∼3μm/pixel for 2X objective and ∼1.5 μm/pixel for 4X objective. To process the data, we developed a MATLAB^®^ pipeline to automatically quantify the root lengths from these experiments. This step is particularly user-friendly since it works without any image preprocessing or human intervention. The platform presented here is highly versatile, and we hope it will be adopted broadly by the scientific community. Keyence^®^ microscopes are affordable compared to confocal microscopes; the imaging chambers used are commercially available and can be used on any microscope types; the script is well-annotated in case users need to finetune several parameters (thresholding value, outliers object size, stitching overlap, distance of the index, stuck /jump threshold) without requiring any coding skills; and the script can work on any type of stitched images, tracking a wide variety of experimental windows ranging from minutes to 12 hours.

We further demonstrated the utility and the power of Root Walker in several case studies. To confirm the sensitivity of the Root Walker platform for capturing early root growth responses, we utilized the exogenous application of IAA and observed a dose-dependent inhibition of root growth just minutes after transfer. At a larger, 12-hour timescale, we investigated the response after transfer using two MAPK inhibitors known to interfere with the MAPK signaling cascade. This revealed that only U0126 triggers a significant root growth decrease, suggesting that MPK2 kinase activity is required to mediate root growth. In the literature, discrepancies between these inhibitors have already been reported in shoots (Bjornson *et al*., 2014). However, opposite to our findings, in shoots, only PD98059 was able to delay the response while no changes were observed in the presence of U0126 highlighting that the sequestration of MPK2 rather than its catalytic activity is critical in triggering this rapid response in shoots. Taken together, this might suggest that there are tissue and time scale dependent MPK2 regulatory mechanisms in plants.

Finally, we have demonstrated that our pipeline is suitable for assessing and harnessing natural variation of root growth and root growth responses at a large scale. The scale of root growth measurements enables genetic screens such as GWAS and subsequent candidate gene identification and validation. Since treatments with chemicals, nutrients, small bioactive molecules, and plant hormones will be easy using this platform, this opens up opportunities to bring about new biological insight using large-scale genetic approaches on fast root growth responses by utilizing natural variation or other collections suitable for genetic discovery (Hauser *et al*., 2013; Coego *et al*., 2014; 1001 Genomes Consortium, 2016; Hu *et al*., 2023). We note that we were able to conduct this experiment in less than 2 weeks with a single person, while in our hands efforts studies that involve the treatment of GWAS accession panels with a single person usually take significantly more time (more than one or two months due to growing plants; re-opening/treating/resealing and scanning plates).

Compared to other publicly available platforms and software, Root Walker provides a simple yet comprehensive, all-in-one, automated solution for the study of early root growth rates (Slovak *et al*., 2014; Durham Brooks, Miller and Spalding, 2010; Betegón-Putze *et al*., 2019; French *et al*., 2009; Geng *et al*., 2013; von Wangenheim *et al*., 2017). We believe that this platform is an excellent choice for conducting large scale screening of early root growth responses to various stimuli, which might enable the identification of the genes and mechanisms involved in root growth regulation. Given its resolution and throughput it might be possible to expand the pipeline to include additional analysis outputs such as kinematic measurements for determining expansion rates of distinct root parts.

## Experimental procedures

### Plant materials and growth conditions

For surface sterilization, *Arabidopsis thaliana* seeds that had been produced under uniform growth conditions and were placed for 1 h in opened 1.5-mL Eppendorf tubes in a sealed box containing chlorine gas generated from 200 mL of 10% sodium hypochlorite and 3.5 mL of 37% hydrochloric acid. Seeds were then put on the surface of media described in Gruber et al., 2013 or ½ MS for GWAS, using 12-cm x 12-cm square plates stratified in the dark at 4 °C for 2-3 days (Gruber *et al*., 2013). The Gruber et al., 2013 media contains, 750 μM of MgSO_4_-7H2O, 625 μM of KH_2_PO_4_, 1000 μM of NH_4_NO_3_, 9400 μM of KNO_3_, 1500 μM of CaCL_2_-2H_2_O, 0.055 μM of CoCL_2_-6H_2_O, 0.053 μM of CuCl_2_-2H_2_O, 50 μM of H_3_BO_3_, 2.5 μM of KI, 50 μM of MnCl_2_-4H_2_O, 0.52 μM of Na_2_MoO_4_-2H_2_O, 15 μM of ZnCl_2_, 75 μM of Na-Fe-EDTA, 1000 μM of MES adjusted to pH 5.5 with KOH and 0.5% sucrose and 1% BactoAgar®. Plants were grown in long day conditions (16/8h) in walk in growth chambers at 21°C, 50uM light intensity, 60% humidity. During nighttime, temperature was decreased to 15°C.

### GWA mapping

260 natural accessions from the Iberic and Swedish subpopulations (12 plants/accession were planted) were grown on 1/2 MS agar plates with 0.5% sucrose and 0.8% of agar and grown for 5 days (16 h light) at 21 °C. About 15 seedlings were then transferred to the chambers (Lab-Tek, Chambers #1.0 Borosilicate Coverglass System, catalog number: 155361) filled with 1/2MS, 0.5% sucrose and 0.8% agar with 100nM EtOH. Note that the transfer took about 45-60 seconds. Images were acquired every minute for 20 minutes, in brightfield conditions using a Keyence^®^ microscope model BZ-X810 with a BZ NIKON Objective Lens (4X) CFI Plan-Apo Lambda. Root image analyses and trait quantification were performed using the MATLAB^®^ script developed in this study “Root_Walker_20min_No_Video_GWAS_Windows.m”(https://github.com/mplatre/Root-Walker-Script.git). Mean, median, variance root growth rate and area under the curve (n ≥ 3 to 15) values were used for GWA study (Table S1). To control for false positives, we used a 5% FDR threshold calculated by the Benjamini–Hochberg–Yekutieli method to correct for multiple testing.

### Phenotyping of the root growth responses IAA and MPK2 inhibitors

Wild type seeds were sowed in media described in Gruber et al., 2013 and stratified in the dark for 2-3 days at 4°C (Gruber *et al*., 2013). Five days after planting, about 20 seedlings were transferred using forceps to an imaging chamber (Lab-Tek, Chambers #1.0 Borosilicate Coverglass System, catalog number: 155361) filled with medium described in Gruber et al., 2013. Note that the transfer took about 45-60 seconds. Images were acquired every minute or 5 minutes 1 hours or 12 hours, respectively, in brightfield conditions using a Keyence^®^ microscope model BZ-X810 with a BZ NIKON Objective Lens (4X or 2X) CFI Plan-Apo Lambda. For manual measurements, time lapse images were used to create videos with the Keyence software^®^. The videos were opened in Fiji (Schindelin *et al*., 2012) and the root tip displacement was measured using the segmented line tool at each time point.

### Phenotyping of root growth responses in plates

Wild type seeds were sowed in media described in Gruber et al., 2013 and stratified in the dark for 2-3 days at 4°C (Gruber *et al*., 2013). Five days after planting, about, 6 plants per genotype were transferred to four12×12-cm plates in a pattern in which the positions of the genotypes were alternating in a block design. After transfer, the plates were scanned every 24 h for 3 days using the BRAT software (ref).

### Treatment with IAA, U0126, PD98059 and PAO

The imaging chamber was filled up with the media described in Gruber et al., 2013 in which IAA dissolved in DMSO (stock solution 1000X compared to the final solution) at 1nM, 10nM or 100nM and DMSO was added (Gruber *et al*., 2013). U0126 and PD98059 dissolved in DMSO (stock solution 20mM or 10mM, respectively) at a concentration of 10μM and DMSO were added to the Gruber media when poured to the imaging chamber. PAO dissolved in DMSO (stock solution 100mM) at a concentration of 7.5, 15 and 30μM and DMSO were added to the gruber media when poured to the imaging chamber.

### Ratio of plasma membrane and cytosol intensity

Confocal images were first denoised using an auto local threshold applying the Otsu method with a radius of 25 and a median filter with a radius of 2 in Fiji(Schindelin *et al*., 2012). To remove every single bright pixel on the generated-binary image the despeckle function was applied. To obtain the plasma membrane skeleton, we detected and removed every intracellular dot using the “Analyze Particles” plugin with the following parameters, size between 0.0001 and 35 000 μm2 and a circularity between 0.18 and 1. Then, we selected and cropped a zone which only showed a proper plasma membrane skeleton. We created a selection from the generated-plasma membrane skeleton and transposed it to the original image. First, we calculated the mean gray value of this zone, corresponding to the plasma membrane mean gray value. Then on the original image the same zone has been removed from and the mean gray value was calculated to obtain the cytosol mean gray value. The plasma membrane mean gray value was then divided by cytosol mean gray value to normalize to obtain the ration of the plasma membrane and the cytosol intensities. This process has been automated in a Macro. An average of 45 cells were used for quantification per root. Every experiment was repeated three times.

### Statistical analysis

Each experiment has been repeated independently at least twice, and in every case the same trend has been recorded for independent experiments, the data have been pooled for further statistical analysis. Each sample was subjected to four different normality tests (Jarque-Bera, Lilliefors, Anderson-Darling and Shapiro-Wilk), and sample were considered as a Gaussian distribution when at least one test was significant (p<0.05) using Xlstat. As a normal distribution was observed a one-way ANOVA coupled with post hoc Tukey or Fisher test was performed (p<0.05) or a two-way student t-test (p<0.05) using Xlstat. As a normal distribution was not observed at two-ways Kruskal-Wallis coupled with post hoc Steel-Dwass-Critchlow-Fligner procedure was performed (p<0.05) using Xlstat. The type of test performed is indicated in the figure legend.

## Acknowledgments

We thank all the Busch lab members for critical discussions, especially Ying Sun. This study was funded by the National Institute of General Medical Sciences of the National Institutes of Health (grant number R01GM127759 to W. Busch), start-up funds from the Salk Institute for Biological Studies and funds from the Salk Harnessing Plants Initiative (W. Busch). M.P. Platre was supported by a long-term postdoctoral fellowship (LT000340/2019 L) by the Human Frontier Science Program Organization.

## Code availability

Matlab^®^ script can be found here, https://github.com/mplatre/Root-Walker-Script.git

## REFERENCES

1001 Genomes Consortium (2016) ‘1,135 Genomes Reveal the Global Pattern of Polymorphism in Arabidopsis thaliana’, Cell, 166(2), pp. 481–491. Available at: 10.1016/j.cell.2016.05.063.

Betegón-Putze, I. et al. (2019) ‘MyROOT: a method and software for the semiautomatic measurement of primary root length in Arabidopsis seedlings’, The Plant Journal, 98(6), pp. 1145–1156. Available at: 10.1111/tpj.14297.

Bjornson, M. et al. (2014) ‘Distinct Roles for Mitogen-Activated Protein Kinase Signaling and CALMODULIN-BINDING TRANSCRIPTIONAL ACTIVATOR3 in Regulating the Peak Time and Amplitude of the Plant General Stress Response1[W][OPEN]’, Plant Physiology, 166(2), pp. 988–996. Available at: 10.1104/pp.114.245944.

Busch, W. et al. (2012) ‘A microfluidic device and computational platform for high-throughput live imaging of gene expression’, Nature Methods, 9(11), pp. 1101–1106. Available at: 10.1038/nmeth.2185.

Coego, A. et al. (2014) ‘The TRANSPLANTA collection of Arabidopsis lines: a resource for functional analysis of transcription factors based on their conditional overexpression’, The Plant Journal: For Cell and Molecular Biology, 77(6), pp. 944–953. Available at: 10.1111/tpj.12443.

Dubey, S.M. et al. (2023) ‘The AFB1 auxin receptor controls the cytoplasmic auxin response pathway in Arabidopsis thaliana’, Molecular Plant, 16(7), pp. 1120–1130. Available at: 10.1016/j.molp.2023.06.008.

Durham Brooks, T.L., Miller, N.D. and Spalding, E.P. (2010) ‘Plasticity of Arabidopsis Root Gravitropism throughout a Multidimensional Condition Space Quantified by Automated Image Analysis’, Plant Physiology, 152(1), pp. 206–216. Available at: 10.1104/pp.109.145292.

Fendrych, M. et al. (2018) ‘Rapid and reversible root growth inhibition by TIR1 auxin signalling’, Nature Plants, 4(7), pp. 453–459. Available at: 10.1038/s41477-018-0190-1.

French, A. et al. (2009) ‘High-Throughput Quantification of Root Growth Using a Novel Image-Analysis Tool’, Plant Physiology, 150(4), pp. 1784–1795. Available at: 10.1104/pp.109.140558.

Ge, X., Fu, Y.M. and Meadows, G.G. (2002) ‘U0126, a mitogen-activated protein kinase kinase inhibitor, inhibits the invasion of human A375 melanoma cells’, Cancer Letters, 179(2), pp. 133–140. Available at: 10.1016/s0304-3835(02)00004-6.

Geng, Y. et al. (2013) ‘A Spatio-Temporal Understanding of Growth Regulation during the Salt Stress Response in Arabidopsis’, The Plant Cell, 25(6), pp. 2132–2154. Available at: 10.1105/tpc.113.112896.

Gomez, R.E. et al. (2022) ‘Phosphatidylinositol-4-phosphate controls autophagosome formation in Arabidopsis thaliana’, Nature Communications, 13(1), p. 4385. Available at: 10.1038/s41467-022-32109-2.

Grossmann, G. et al. (2011) ‘The RootChip: An Integrated Microfluidic Chip for Plant Science’, THE PLANT CELL ONLINE, 23(12), pp. 4234–4240. Available at: 10.1105/tpc.111.092577.

Gruber, B.D. et al. (2013) ‘Plasticity of the Arabidopsis Root System under Nutrient Deficiencies’, Plant Physiology, 163(1), pp. 161–179. Available at: 10.1104/pp.113.218453.

Hauser, F. et al. (2013) ‘A Genomic-Scale Artificial MicroRNA Library as a Tool to Investigate the Functionally Redundant Gene Space in Arabidopsis[W]’, The Plant Cell, 25(8), pp. 2848–2863. Available at: 10.1105/tpc.113.112805.

Hotokezaka, H. et al. (2002) ‘U0126 and PD98059, specific inhibitors of MEK, accelerate differentiation of RAW264.7 cells into osteoclast-like cells’, The Journal of Biological Chemistry, 277(49), pp. 47366–47372. Available at: 10.1074/jbc.M208284200.

Hsu, C.-W. et al. (2022) ‘Protocol for fast scRNA-seq raw data processing using scKB and non-arbitrary quality control with COPILOT’, STAR Protocols, 3(4), p. 101729. Available at: 10.1016/j.xpro.2022.101729.

Hu, Y. et al. (2023) ‘Multi-Knock—a multi-targeted genome-scale CRISPR toolbox to overcome functional redundancy in plants’, Nature Plants, 9(4), pp. 572–587. Available at: 10.1038/s41477-023-01374-4.

Jia, Z., Giehl, R.F.H. and von Wirén, N. (2022) ‘Nutrient–hormone relations: Driving root plasticity in plants’, Molecular Plant, 15(1), pp. 86–103. Available at: 10.1016/j.molp.2021.12.004.

Li, B., Sun, L., Huang, J., Goschl, C., et al. (2019) ‘GSNOR provides plant tolerance to iron toxicity viapreventing iron-dependent nitrosative and oxidative cytotoxicity’, Nature Communications, 10, p. 13.

Li, B., Sun, L., Huang, J., Göschl, C., et al. (2019) ‘GSNOR provides plant tolerance to iron toxicity via preventing iron-dependent nitrosative and oxidative cytotoxicity’, Nature Communications, 10(1), p. 3896. Available at: 10.1038/s41467-019-11892-5.

Li, L. et al. (2021) ‘Antagonistic cell surface and intracellular auxin signalling regulate plasma membrane H+-fluxes for root growth’, Nature, 599(7884), pp. 273–277. Available at: 10.1038/s41586-021-04037-6.

de Luis Balaguer, M.A. et al. (2016) ‘Multi-sample Arabidopsis Growth and Imaging Chamber (MAGIC) for long term imaging in the ZEISS Lightsheet Z.1’, Developmental Biology, 419(1), pp. 19–25. Available at: 10.1016/j.ydbio.2016.05.029.

Morris, E.C. et al. (2017) ‘Shaping 3D Root System Architecture’, Current Biology, 27(17), pp. R919–R930. Available at: 10.1016/j.cub.2017.06.043.

Noack, L.C. et al. (2021) ‘A nanodomain-anchored scaffolding complex is required for the function and localization of phosphatidylinositol 4-kinase alpha in plants’, The Plant Cell, 34(1), pp. 302–332. Available at: 10.1093/plcell/koab135.

Ogura, T. et al. (2019) ‘Root System Depth in Arabidopsis Is Shaped by EXOCYST70A3 via the Dynamic Modulation of Auxin Transport’, Cell, 178(2), pp. 400–412.e16. Available at: 10.1016/j.cell.2019.06.021.

Okazaki, K., Miyagishima, S. and Wada, H. (2015) ‘Phosphatidylinositol 4-Phosphate Negatively Regulates Chloroplast Division in Arabidopsis’, The Plant Cell Online, 27(3), pp. 663–674. Available at: 10.1105/tpc.115.136234.

Platre, M.P. et al. (2018) ‘A Combinatorial Lipid Code Shapes the Electrostatic Landscape of Plant Endomembranes’, Developmental Cell, 45(4), pp. 465–480.e11. Available at: 10.1016/j.devcel.2018.04.011.

Platre, M.P. et al. (2022) ‘The receptor kinase SRF3 coordinates iron-level and flagellin dependent defense and growth responses in plants’, Nature Communications, 13(1), p. 4445. Available at: 10.1038/s41467-022-32167-6.

Prigge, M.J. et al. (2020) ‘Genetic analysis of the Arabidopsis TIR1/ AFB auxin receptors reveals both overlapping and specialized functions’, eLife, 9(54740), p. 28.

Satbhai, S.B. et al. (2017) ‘Natural allelic variation of FRO2 modulates Arabidopsis root growth under iron deficiency’, Nature Communications, 8, p. 15603. Available at: 10.1038/ncomms15603.

Schindelin, J. et al. (2012) ‘Fiji: an open-source platform for biological-image analysis’, Nature Methods, 9(7), pp. 676–682. Available at: 10.1038/nmeth.2019.

Serre, N.B.C. et al. (2021) ‘AFB1 controls rapid auxin signalling through membrane depolarization in Arabidopsis thaliana root’, Nature Plants, 7(9), pp. 1229–1238. Available at: 10.1038/s41477-021-00969-z.

Shih, H.-W. et al. (2015) ‘The Cyclic Nucleotide-Gated Channel CNGC14 Regulates Root Gravitropism in Arabidopsis thaliana’, Current Biology, 25(23), pp. 3119–3125. Available at: 10.1016/j.cub.2015.10.025.

Simon, M.L.A. et al. (2016) ‘A PtdIns(4)P-driven electrostatic field controls cell membrane identity and signalling in plants’, Nature Plants, 2(7), p. nplants201689. Available at: 10.1038/nplants.2016.89.

Slovak, R. et al. (2014) ‘A Scalable Open-Source Pipeline for Large-Scale Root Phenotyping of Arabidopsis’, The Plant Cell, 26(6), pp. 2390–2403. Available at: 10.1105/tpc.114.124032.

Slovak, R. et al. (2020) ‘Ribosome assembly factor Adenylate Kinase 6 maintains cell proliferation and cell size homeostasis during root growth’, The New Phytologist, 225(5), pp. 2064–2076. Available at: 10.1111/nph.16291.

Stanley, C.E. et al. (2018) ‘Dual-flow-RootChip reveals local adaptations of roots towards environmental asymmetry at the physiological and genetic levels’, The New Phytologist, 217(3), pp. 1357–1369. Available at: 10.1111/nph.14887.

Taylor, I. et al. (2021) ‘Mechanism and function of root circumnutation’, Proceedings of the National Academy of Sciences, 118(8), p. e2018940118. Available at: 10.1073/pnas.2018940118.

Wang, L. et al. (2018) ‘Negative regulator of E2F transcription factors links cell cycle checkpoint and DNA damage repair’, Proceedings of the National Academy of Sciences, 115(16), pp. E3837–E3845. Available at: 10.1073/pnas.1720094115.

von Wangenheim, D. et al./person-group>. (2017) ‘Live tracking of moving samples in confocal microscopy for vertically grown roots’, eLife. Edited by C.S. Hardtke, 6, p. e26792. Available at: 10.7554/eLife.26792.

Xiong, Y. et al. (2013) ‘Glucose–TOR signalling reprograms the transcriptome and activates meristems’, Nature, 496(7444), pp. 181–186. Available at: 10.1038/nature12030.

Xiong, Y. and Sheen, J. (2012) ‘Rapamycin and Glucose-Target of Rapamycin (TOR) Protein Signaling in Plants*’, Journal of Biological Chemistry, 287(4), pp. 2836–2842. Available at: 10.1074/jbc.M111.300749.

