## Supplementary Figures for "Root Walker: an automated pipeline for large scale quantification of early root growth responses at high spatial and temporal resolution"

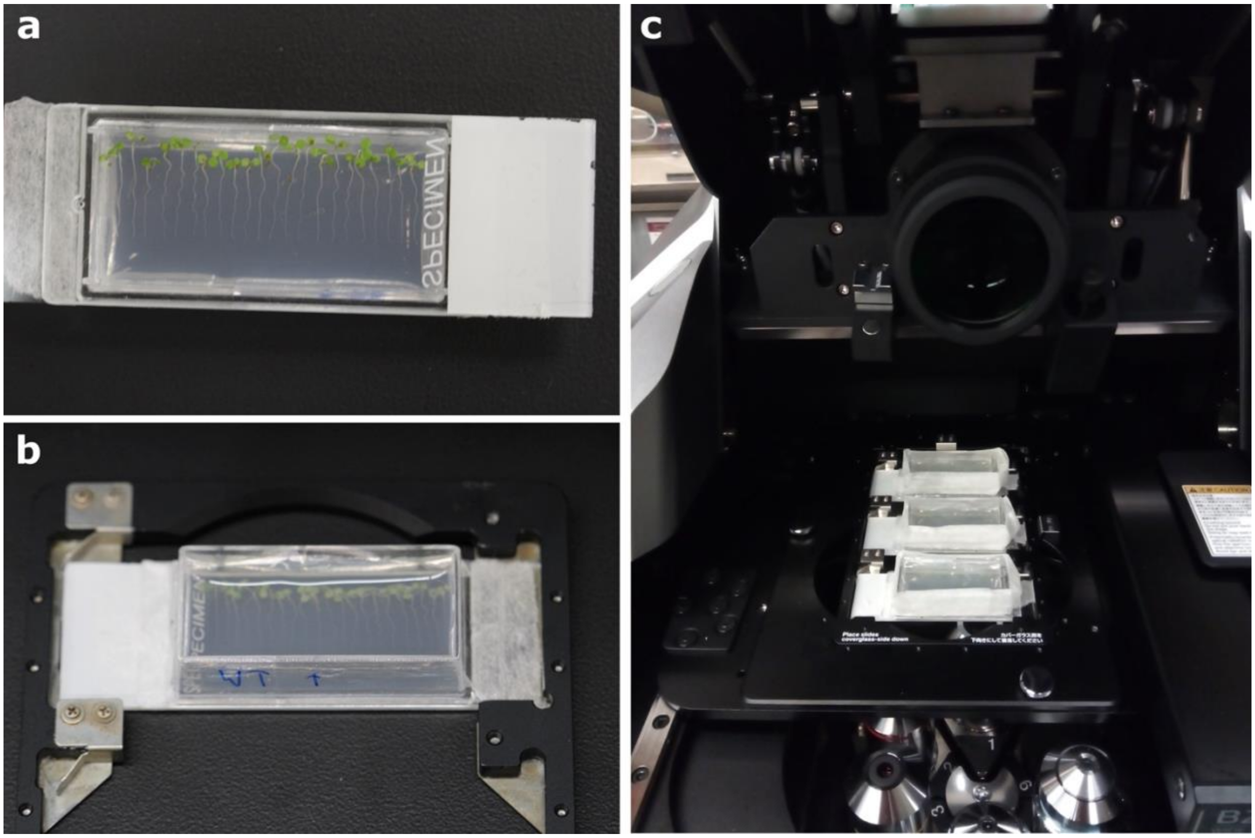


**Figure S1. Microscope and chamber set up**. (**a**) Chamber assembly for root growth time lapse imaging. (**b**) Picture of the chamber loaded on the stage for imaging for 60 minutes. (**c**) picture of the three chambers loaded on the microscope allowing multiple imaging conditions and genotypes for 12 hours.

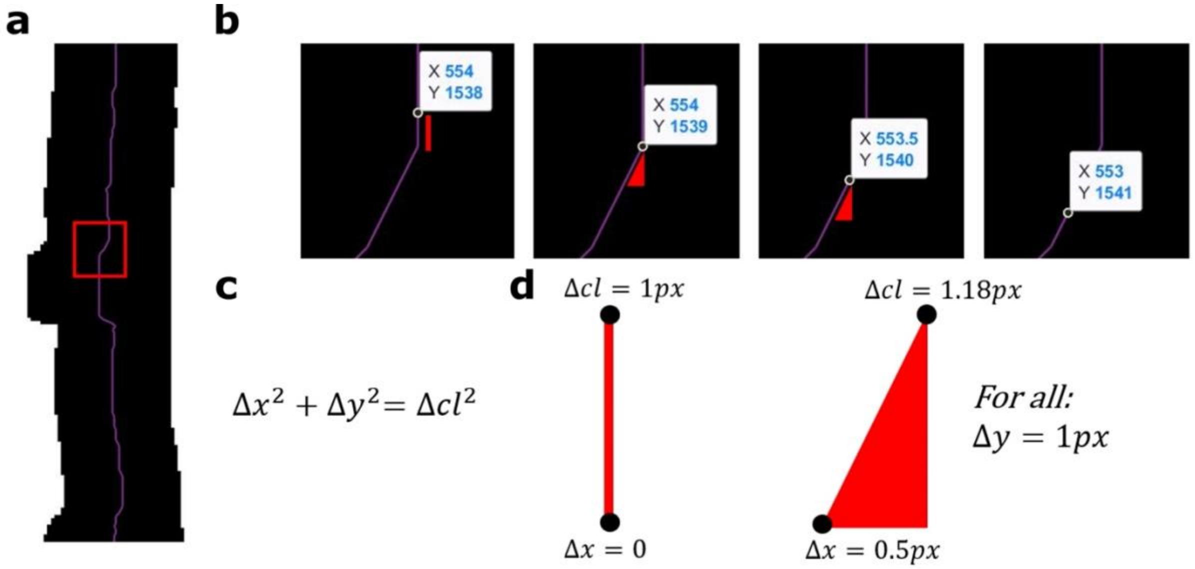


**Figure S2. Center line length calculation methodology**. (**a**) Image showing a part of a segmented root. Black pixels: Segmented root; White pixels: background; Purple pixels: Center line. (**b**) Coordinates (X, Y) of the root center array along the center line, when walking the root center line for one pixel in the y-direction. (**c**) Equation to calculate the root growth and subsequent root length at each time step in pixel. (**d**) Example of length calculations for straight and angled lines. cl, length in pixel; px, pixel.

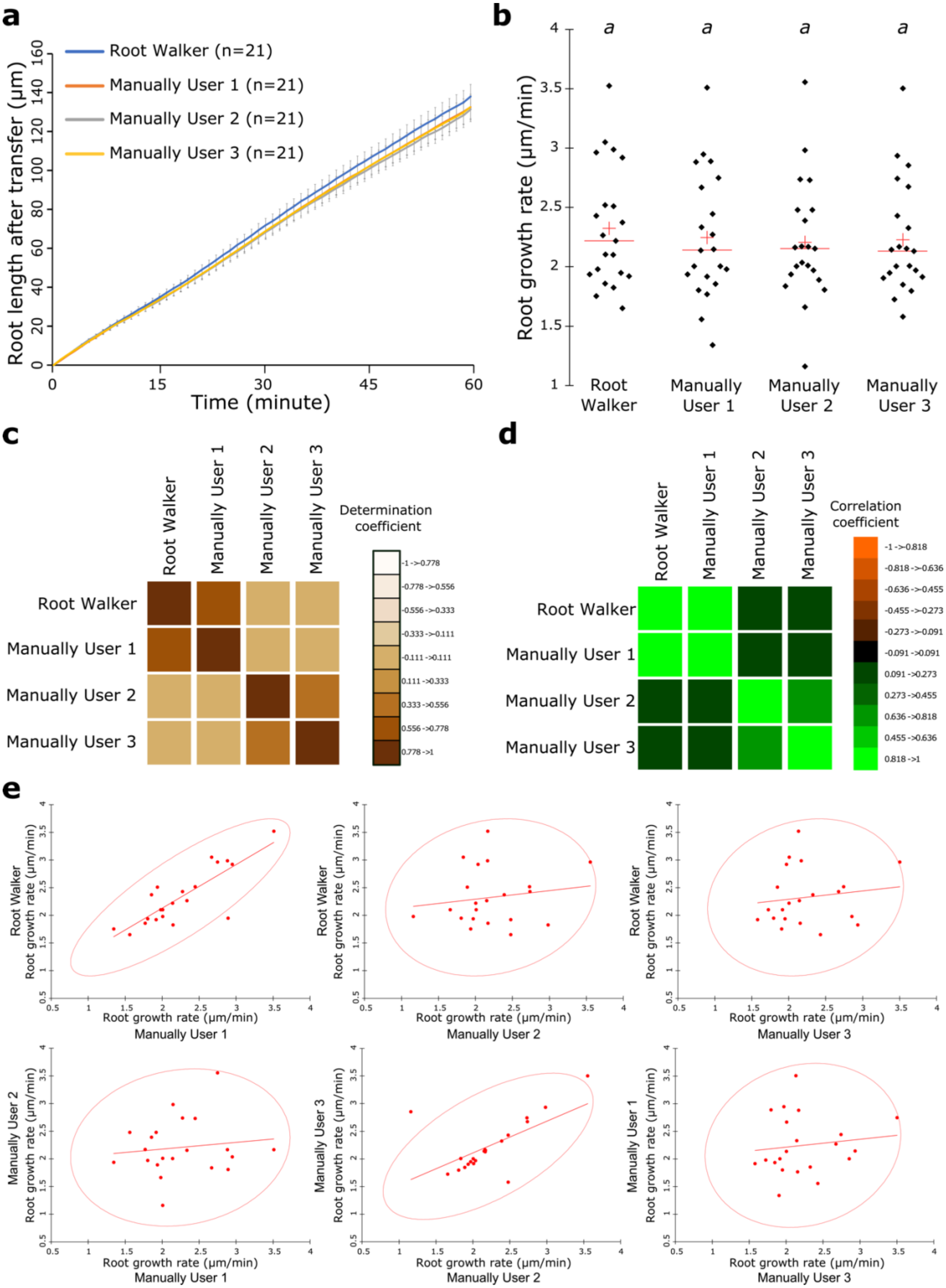
a
Root Walker (n=21)
Manually User 1 (n=21)
Manually User 2 (n=21)
Manually User 3 (n=21)
Root length after transfer (µm)
Time (minute)
b
a
Root growth rate (µm/min)
Root Walker
Manually User 1
Manually User 2
Manually User 3
c
Root Walker
Manually User 1
Manually User 2
Manually User 3
Determination coefficient
-1 ->0.778
-0.778 ->0.556
-0.556 ->0.333
-0.333 ->0.111
-0.111 ->0.111
0.111 ->0.333
0.333 ->0.556
0.556 ->0.778
0.778 ->1
d
Root Walker
Manually User 1
Manually User 2
Manually User 3
Correlation coefficient
-1 ->0.818
-0.818 ->0.636
-0.636 ->0.455
-0.455 ->0.273
-0.273 ->0.091
-0.091 ->0.091
0.091 ->0.273
0.273 ->0.455
0.636 ->0.818
0.455 ->0.636
0.818 ->1
e
Root Walker Root growth rate (µm/min)
Root growth rate (µm/min) Manually User 1
Root growth rate (µm/min) Manually User 2
Root growth rate (µm/min) Manually User 3
Manually User 2 Root growth rate (µm/min)
Manually User 3 Root growth rate (µm/min)
Manually User 1 Root growth rate (µm/min)

**Figure S3. Comparison of root growth rate between Root Walker analysis script and human observer.** (**a**) Average root length of 7-day old seedlings after transfer to the imaging chamber containing standard medium and quantified by the Root Walker script or manually by user 1, 2 and 3. Error bars represent the standard deviation. (**b**) Root growth rate of data presented in a. [[one-way ANOVA followed by a Fisher LSD test; letters indicate statistical differences ($p<0.05$)]. n.s., non-significant. Red crosses represent the average and the red bars the median. Black dots individual measurements. (**c**) Image of the matrix of coefficient of determination. (**d**) Image of the correlation matrix (Pearson coefficient). (**e**) Scatter plots of measured values by user 1,2, and 3 and Root Walker. Red circle indicates a confidence interval (95%) according to Fisher procedure.

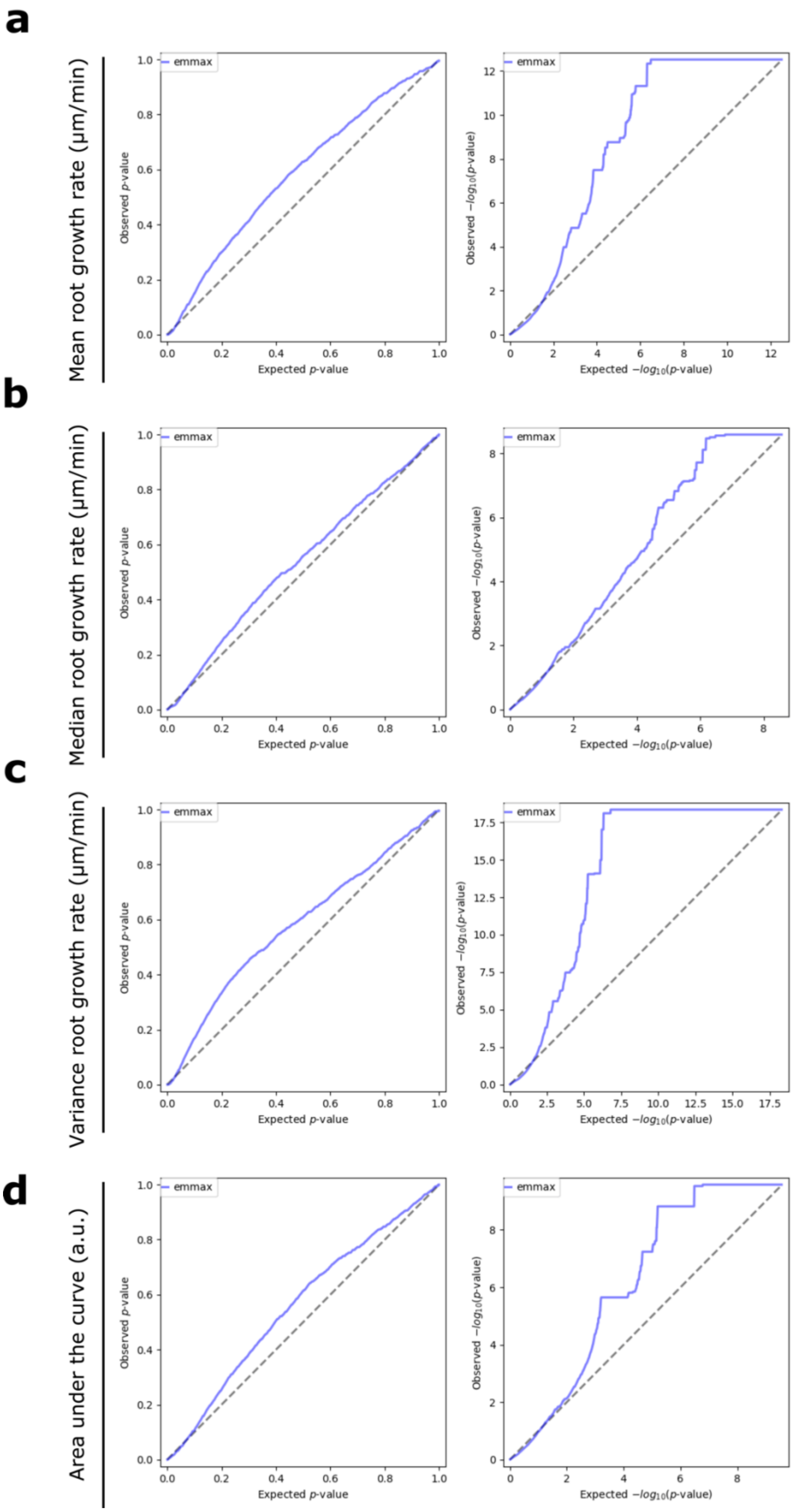
a
Mean root growth rate (µm/min)
b
Median root growth rate (µm/min)
c
Variance root growth rate (µm/min)
d
Area under the curve (a.u.)
emmax
Observed p-value
Expected p-value
Observed −log10(p-value)
Expected −log10(p-value)

**Figure S4. Q-Q plot related to GWAS**. (**a-d**), Q-Q plot of mean root growth rate (**a**), median root growth rate (**b**), variance root growth rate (**c**), area under the curve (**d**).

**Video S1. Output video after root length calculation.** Representative video generated by the root Walker script. Upper, raw video of all the roots detected and depicted by the root center line in color. R represents the number of roots detected. Lower, binary video of all the roots which have passed the “jump” and “stuck” threshold.

**Video S2. Video of roots grown under different concentration of IAA for an hour.** From left to right, representative video of roots of 5-day old seedlings treated with DMSO, IAA at 1nM, 10nM and 100nM for an hour. Scale bar, 100 µm.

**Video S3. Video of roots grown under different MPK inhibitors for 12 hours.** From left to right, representative video of roots of 5-day old seedlings treated with DMSO, PD98059 and U0126 at 10µM for 12 hours. Scale bar, 200 µm.

**Video S4. Video of roots grown under different concentration of PAO for 20 minutes.** From left to right, representative video of roots of 5-day old seedlings treated with DMSO, PAO 7.5µM, PAO 15µM and PAO 30µM over the course of 20 minutes. Scale bar, 100 µm.

**Video S5. Video of roots grown under different concentration of IAA for an hour.** From left to right, video of 5 to 8-day old Arabidopsis seedlings in DMSO (left) and β-estradiol (β-EST, right) at 2.5µM of WT, TOR-ES#1, TOR-ES#2, ERF-1#1, ERF-1#2, ERF-1#3. Scale bar, 1 cm.
